# A single cell transcriptomic analysis of human neocortical development

**DOI:** 10.1101/401885

**Authors:** Damon Polioudakis, Luis de la Torre-Ubieta, Justin Langerman, Andrew G. Elkins, Jason L. Stein, Celine K. Vuong, Carli K. Opland, Daning Lu, William Connell, Elizabeth K. Ruzzo, Jennifer K. Lowe, Tarik Hadzic, Flora I. Hinz, Shan Sabri, William E. Lowry, Kathrin Plath, Daniel H. Geschwind

## Abstract

Defining the number, proportion, or lineage of distinct cell types in the developing human brain is an important goal of modern brain research. We defined single cell transcriptomic profiles for 40,000 cells at mid-gestation to identify cell types in the developing human neocortex. We define expression profiles corresponding to all known major cell types at this developmental period and identify multiple transcription factors and co-factors expressed in specific cell types, providing an unprecedented resource for understanding human neocortical development including the first single-cell characterization of human subplate neurons. We characterize major developmental trajectories during early neurogenesis, showing that cell type differentiation occurs on a continuum that involves transitions that tie cell cycle progression with early cell fate decisions. We use these data to deconvolute regulatory networks and map neuropsychiatric disease genes to specific cell types, implicating dysregulation of specific cell types, as the mechanistic underpinnings of several neurodevelopmental disorders. Together these results provide an extensive catalog of cell types in human neocortex and extend our understanding of early cortical development, human brain evolution and the cellular basis of neuropsychiatric disease.

**One Sentence Summary:** Comprehensive single cell transcriptomes in developing human cortex inform models of cell diversity, differentiation and disease risk.

## Main text

The human cortex is composed of billions of cells estimated to encompass hundreds or thousands of distinct cell types each with unique functions (*1, 2*). It is reasonable to assume that to understand a system as complex as the human cortex, it is necessary to understand its components (*3, 4*). Ground breaking work in mouse revealed the power of single-cell transcriptomics for providing a framework for understanding the complexity and heterogeneity of cell types in the brain (*5-10*). The availability of high quality tissue and advances in single-cell transcriptomic technologies now permit us to catalog the cell type diversity of the human cortex in a comprehensive and unbiased manner (*11*). Despite the enormous progress that has been made in characterizing early cortical development (*1, 12-16*), many of the molecular mechanisms underpinning the generation, differentiation, and development of the diverse types of cells remain largely unknown (*17*). Molecular taxonomies of cortical cell types from developing human brains enable us to understand the mechanisms of neurogenesis and how the remarkable cellular diversity found in the human cortex is achieved (*18-22*). Developing knowledge of neocortical cell types and their transcriptional programs during this epoch is a key step in understanding mechanistic dysregulation in neurodevelopmental disorders with cell type resolution. Several recent studies have taken a first step in this direction, analyzing several hundred, or a few thousand cells from developing human brain (*18-22*). However, advances in technology and throughput [*e.g.* Drop-seq (*5*)] now allow us to analyze an order of magnitude more cells to complement and extend these studies, providing a deeper picture of human cortical development.

Here we use single-cell RNA sequencing (scRNA-seq) to define cell types and compile cell type transcriptomes in the developing human neocortex. We focus on the cortical anlage at mid-gestation (gestation week (GW) 17 to 18) (**Fig. 1A**) because this period contains the major germinal zones and the developing cortical laminae containing migrating and newly born neurons, and neurodevelopmental processes occurring during this epoch are implicated in neuropsychiatric disease (*23, 24*). To optimize detection of distinct cell types, we separated the cortex into the germinal zones [ventricular zone (VZ) and subventricular zone (SVZ)] and developing cortex [subplate (SP) and cortical plate (CP)] prior to single-cell isolation. Using Drop-seq (*5*), we obtained and compared high quality profiles for ∼40,000 cells from human cortex (**Fig. 1A**; fig. S1, A and B; and tables S1 to S3), and a small subset with microfluidics approaches (Fluidigm) for technical comparisons.

**Fig. 1.**
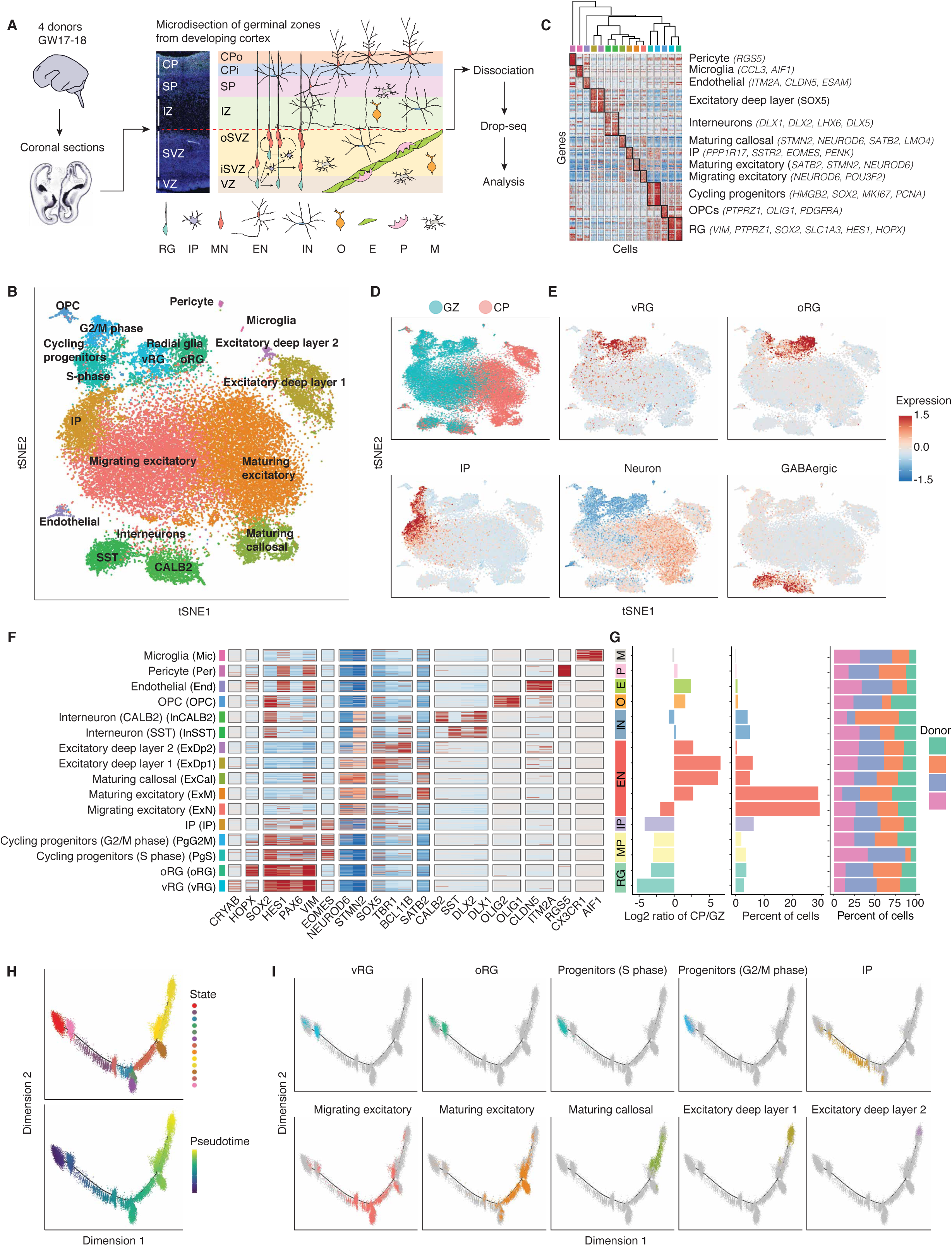
A catalog of cell types in developing human neocortex.(A) Diagram of experimental design illustrating anatomical dissection of regions. VZ: ventricular zone; iSVZ: inner subventricular zone; oSVZ: outer subventricular zone; IZ: intermediate zone; SP: subplate; CPi: inner cortical plate; CPo: outer cortical plate; RG: radial glia; IP: intermediate progenitor; MN: newborn migrating excitatory neuron; EN: excitatory neuron; IN: interneuron; O: oligodendrocyte precursor; E: endothelial cell; P: pericyte; M: microglia. (B) Scatter plot visualization of cells after principal components analysis and t-stochastic neighbor embedding (tSNE), colored by Seurat clustering, and annotated by major cell types. (C) Heatmap of gene expression for each cell. Cells are grouped by Seurat clustering, and the mean expression profile of enriched genes for each cluster was used to hierarchically cluster the Seurat clusters. The top 20 most enriched genes are shown per cluster, and anatomical marker genes in the top 20 most enriched genes [for each cluster] are noted in parentheses. Color bar matches Seurat clusters in B. (D and E) tSNE of cells colored by anatomical source (D), or mean expression of groups of canonical marker genes of major cell types (E). (F) Heatmap of expression profiles of canonical cell type marker genes. Cells are grouped by Seurat clustering. Color bar matches Seurat clusters in B. (G) Cluster metrics. Ratio of cells derived from GZ or CP. Percent of total cell population. Percent of cells derived from each donor. Bar colors indicate grouping of cells by major cell type, *e.g.* CALB2 and SST interneurons are both blue. (H) Pseudo-time analysis using Monocle 2.0 of cells expected to be part of the neurogenesis-differentiation axis, colored by Monocle state or pseudo-time. Each point represents a cell. Monocle 2.0 is a computational method aimed at performing lineage trajectory reconstruction based on single-cell transcriptomics data. Pseudo-time represents an ordering of cells based upon the inferred trajectory, or in other words the predicted lineage trajectory. (I) Pseudo-time trajectory colored by Seurat clusters.

We first applied standard unbiased clustering based on stochastic nearest neighbor imbedding (tSNE; see materials and methods) and spectral K-nearest neighbor graph based clustering (*25*), identifying 16 transcriptionally distinct cell groups. Cell types originated from the expected anatomical source, and clustered by biological cell type rather than batch or technical artifacts (**Fig. 1, B to G** and fig. S1, C and D). We identified multiple groups of cells at different stages of neuronal differentiation and maturation, corresponding to all known major cell types at this developmental time period including: radial glia (RG), intermediate progenitors (IP), migrating, SP and CP excitatory neurons (Ex), interneurons (In), microglia (Mic), oligodendrocyte precursors (OPC), and supporting cells (endothelial cells, pericytes) (**Fig. 1, B to F;** fig. S2A; and tables S4 to S6). Clusters contained between 50 and 2,000 cells, with the smallest cluster captured, which belonged to microglia, comprised of ∼50 cells, with other small clusters for oligodendrocyte precursors, endothelia, and pericytes comprised of 306, 237, 114 cells, respectively (**Fig. 1G** and table S4). Clusters were reproducible and robust as ascertained by bootstrapping (fig. S2B). Ordering of cells by pseudo-time in an unbiased manner using Monocle 2, a computational method that performs lineage trajectory reconstruction based on single-cell transcriptomics data, (*26, 27*) confirmed the predicted developmental trajectory (**Fig. 1, H and I**). For example, it is possible observe the ordered transitions between different neural progenitor types and maturing glutamatergic neurons, with RG transitioning to IPs, and IPs transitioning to newborn migrating neurons (**Fig. 1I**).

Recognizing that with scRNA-seq there is an inherent practical tension between sequencing depth and the number of cells profiled, we explored the consequences of this tradeoff by performing down-sampling of sequencing depth and the number of cells analyzed to determine thresholds for cell type detection (fig. S3, A to D). We observed that cell type detection, especially for low abundance types appears to be more sensitive to the number of cells profiled than sequencing depth (fig. S3, A to D). This is also consistent with the observation that among the genes most highly expressed in a given cluster, most are specific to a cell type, and not simply highly expressed house-keeping genes (fig. S4, A and B). Further, while each individual cell profile is an incomplete representation of that cell type (*28*), we reasoned that we could harness the power of sampling single cells at a large scale by pooling transcriptomes within cells of a given type to provide more complete cell type transcriptome representations. To evaluate the completeness of cell type signatures derived from pooling, we iteratively subsampled cells from clusters and compared the stability of gene expression signatures across iterations by ranking genes by mean expression level (figs. S4, C and D). Subsamples of 50 cells showed high stability in ranking (see materials and methods) for the top 1,000 genes, whereas 500 cells showed very high rank stability for the top 4,000 genes, allowing us to empirically assess the completeness of cell type signatures with different sample sizes (figs. S4, C and D). At a depth of 40,000 cells, we obtain stable transcriptomes representing 3,000-5,000 genes for most of the cell types present (table S5). Comparison of these data with a lower throughput, higher sequencing depth method (Fluidigm C1, (*19*), fig. S5), revealed that although Fluidigm C1 detected a greater number of genes on average from individual cells, the ability to leverage an order of magnitude more cells with Drop-seq provided more stable mRNA transcript profiles for a given cell type (fig. S5, A to C). We provide these cell type specific expression profiles with annotated gene expression ranking confidence measures for each cell type (table S5).

To further evaluate the quality of scRNA-seq transcriptomic catalogs, we compared scRNA-seq datasets produced from different technologies and laboratories to bulk RNA-seq expression profiles from human fetal cortex (*29*). We reasoned that pooling scRNA-seq expression profiles from many cells from a tissue would reveal the magnitude to which single cell protein-coding transcriptomes approximate bulk tissue expression profiles. Consistently, gene expression profiles generated using different scRNA-seq methodologies strongly correlated with bulk RNA-seq gene expression profiles (Spearman 0.69-0.83) (fig. S6, A and B). However, we did observe a group of approximately 400 protein coding genes that were consistently under-represented in single-cell datasets across the methods and laboratories compared to bulk tissue RNA-seq (fig. S6, C and D). These genes encoded cell adhesion molecules, were significantly longer, brain enriched, and involved in neuronal development (fig. S6, E to H). The differences between scRNA-seq methods and bulk tissue RNA-seq may be due to mapping differences between poly-A primed scRNA-seq datasets and rRNA depleted full length mRNA bulk tissue RNA-seq, or dissociation procedures common to all scRNA-seq methods. However, comparison of expression of canonical cell type marker genes showed similar expression levels compared to bulk tissue RNA-seq (fig. S6I). This indicates that despite small biases in gene detection shared across scRNA-seq methods, the relative frequencies of major cell types were not over- or under-represented in the Drop-seq dataset. In conjunction with the high correlation with bulk tissue RNA-seq this demonstrates the reproducibility and robustness of the scRNA-seq dataset (fig. S6, A and I).

Having ascertained the quality and robustness of the cell clusters and associated gene expression profiles, we reasoned that we could gain insight into cell type specific regulatory programs by comparing transcription factor expression across cell types. Indeed, we find previously characterized transcription factors and co-factors enriched in their corresponding cell types (**Figs. 1F and 2A**). In addition, we find several transcription factors and co-factors (ZFHX4, CARHSP1, ST18, and CSRP2) that have not been associated with specific neocortical cell types (**Fig. 2A).** These transcription factors also displayed laminae-specific expression in a bulk tissue laser captured micro-dissected (LCM) expression dataset (*30*), and temporal trajectories similar to canonical cell type markers (**Fig. 2, A and B** and fig. S7, A and B).

**Fig. 2.**
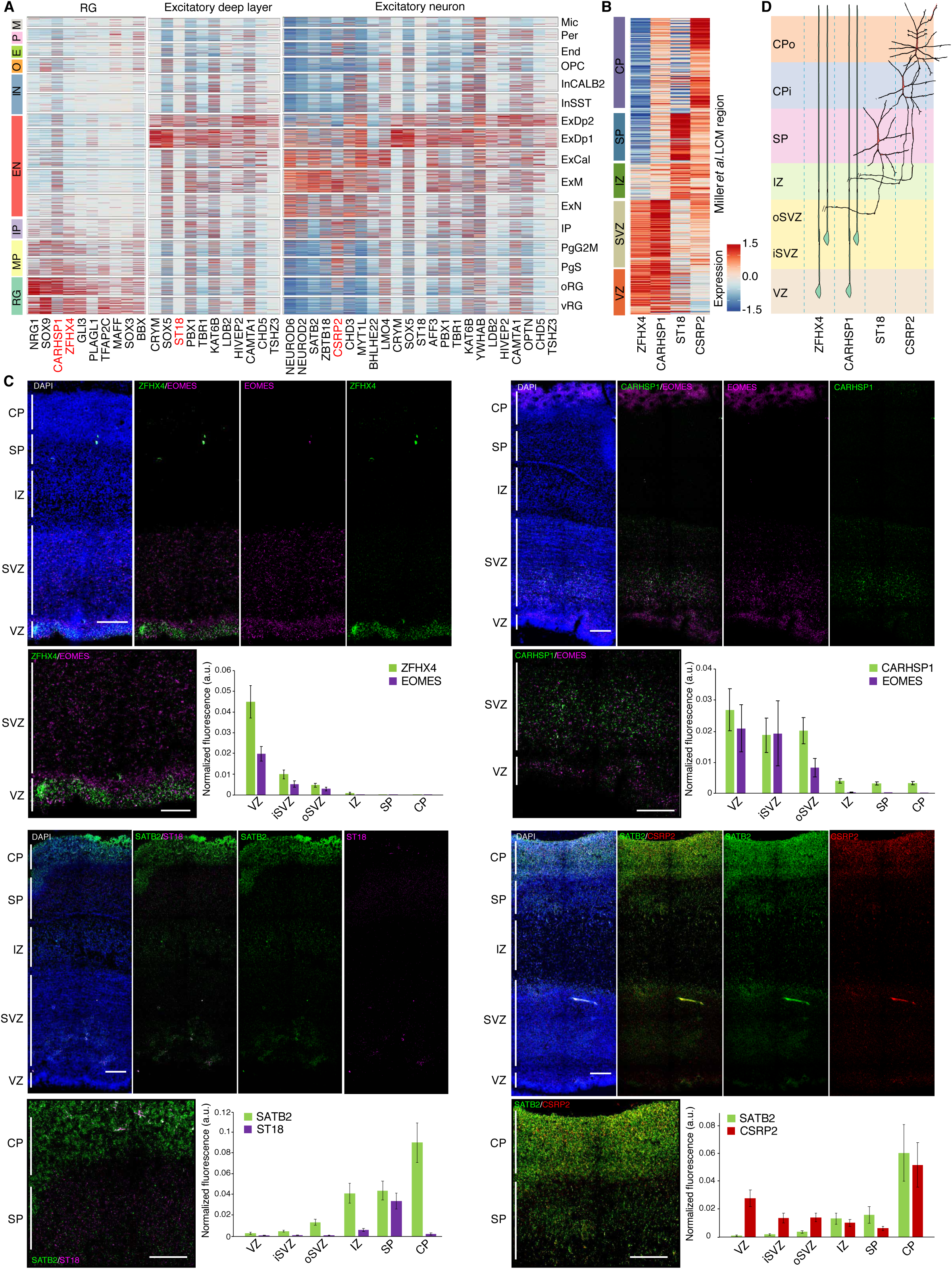
Novel cell type enrichment of transcription factors and co-factors. (A) Heatmap of expression of transcription factors, co-factors, and chromatin remodelers enriched in RG, excitatory neurons, and deep layer excitatory neurons. Cells are grouped by cluster. Red indicates factors previously unknown to be enriched in the neocortical cell types of interest. (B) Expression of factors of interest in bulk tissue LCM laminae from developing cortex. (C) RNA FISH of fetal cortex probed with the newly identified cell-enriched transcription factors CARHSP1 and ZFHX4 (RG in the VZ and SVZ), ST18 (SP neurons), and CSRP2 (glutamatergic neurons in the CP). Quantification of normalized fluorescence intensity per layer for each set of probes (see materials and methods). Scale bar = 250μm (left) or 100μm (inset). (D) Schematic of cell type specific expression of factors of interest. RG: radial glia; MP: mitotic progenitor; IP: intermediate progenitor; EN: excitatory neuron; IN: interneuron; O: oligodendrocyte precursor; E: endothelial cell; P: pericyte; M: microglia; VZ: ventricular zone; iSVZ: inner subventricular zone; oSVZ: outer subventricular zone; IZ: intermediate zone; SP: subplate; CPi: inner cortical plate; CPo: outer cortical plate

The ST18-expressing cluster showed a division in cells expressing deep layer markers and cells expressing SP markers (fig. S8, A and B) (*31, 32*). However, SP markers previously defined in other species were also expressed in subsets of upper layer neurons and neural progenitors (fig. S8B) and were not uniquely expressed in the SP in a fetal LCM atlas (*30*) (fig. S8C), possibly due to the temporal specificity of many of these markers (*31*). To derive a human SP set of markers across mid-gestation, we used the LCM dataset to identify SP enriched genes (fig. S8C, see materials and methods). This new group of SP markers displayed clear expression in a group of cells within the deep layer excitatory cluster, overlapping with ST18 (fig. S8B); sub-clustering (see materials and methods) separated the deep layer neurons from the SP neurons (fig. S8D). Genes enriched in the SP neuron cluster, or highly correlated with ST18 display SP enrichment in the LCM laminae dataset, verifying our capture of SP neurons and identification of many additional SP neuron markers (fig. S8, E to H). This represents the first transcriptomic characterization of SP neurons at single-cell resolution, and a valuable resource for exploring SP biology.

To validate predictions for these putative novel cell type markers, we performed RNA FISH, which confirmed laminae-specific expression of each of the transcription factors and co-factors tested: *ZFHX4* and *CARHSP1* in RG, *ST18* in the SP, and *CSRP2* in excitatory neurons (**Fig. 2, C and D**). Of particular interest was *ZFHX4*, which has been previously associated with 8q21.11 microdeletion, a syndrome characterized by intellectual disability, hypotonia, decreased balance, sensorineural hearing loss, and unusual behavior (*33*). Our data localizes ZFHX4 specifically to RG in the developing human neocortex for the first time, implicating specific dysregulation of RG as the mechanism underlying 8q21.11 microdeletion syndrome.

A first step in understanding the regulatory mechanisms active in the human neocortex is assigning the activity of regulatory elements to specific cell types. To deconvolute the cell type specificity of regulatory elements in human fetal cortex we leveraged a recently generated map of regulatory elements active in developing fetal cortex and their putative target genes (*29*). Using this map, we identify promoters and enhancers regulating the expression of genes enriched in each of the cell clusters defined in this study (see materials and methods) (table S7). Enhancers associated with specific cell types were characterized by remarkable consistency in mean enhancer size, number associated with each gene, and distance to the target gene for each cell type (fig. S9, A to F). In addition, there was no correlation between target gene length or GC content and number of associated enhancers (fig. S9, G and H). This represents the first map of active regulatory elements regulating cell-type specific genes in the human neocortex and a rich resource for understanding human neocortical regulatory mechanisms.

Neurons are generated from the controlled asymmetric division of neural progenitors, which prompted us to analyze the distinct transcriptional states of cycling cells during this process (*16*). Neural progenitors clustered by cell cycle state in addition to cell type (**Figs. 1E and 3A**, and fig S10, A to C), with about 30% of progenitors cycling, roughly consistent with previous observations (37% based on immunostaining) (*34*). Remarkably, we also observed that many of the cycling progenitors individually expressed markers of several distinct major cell types, including RGs, IPs, and neurons (**Fig. 3, A to C**). Doublets were an insufficient explanation for the co-expression of distinct cell type makers for multiple reasons, including that the number of cells expressing multiple major cell type markers is twice the empirically assessed doublet rate (table S2 and fig. S1B—also see materials and methods) and the highly non-random distribution of the cell types expressing markers of two cell types (**Fig. 3C**).

**Fig. 3.**
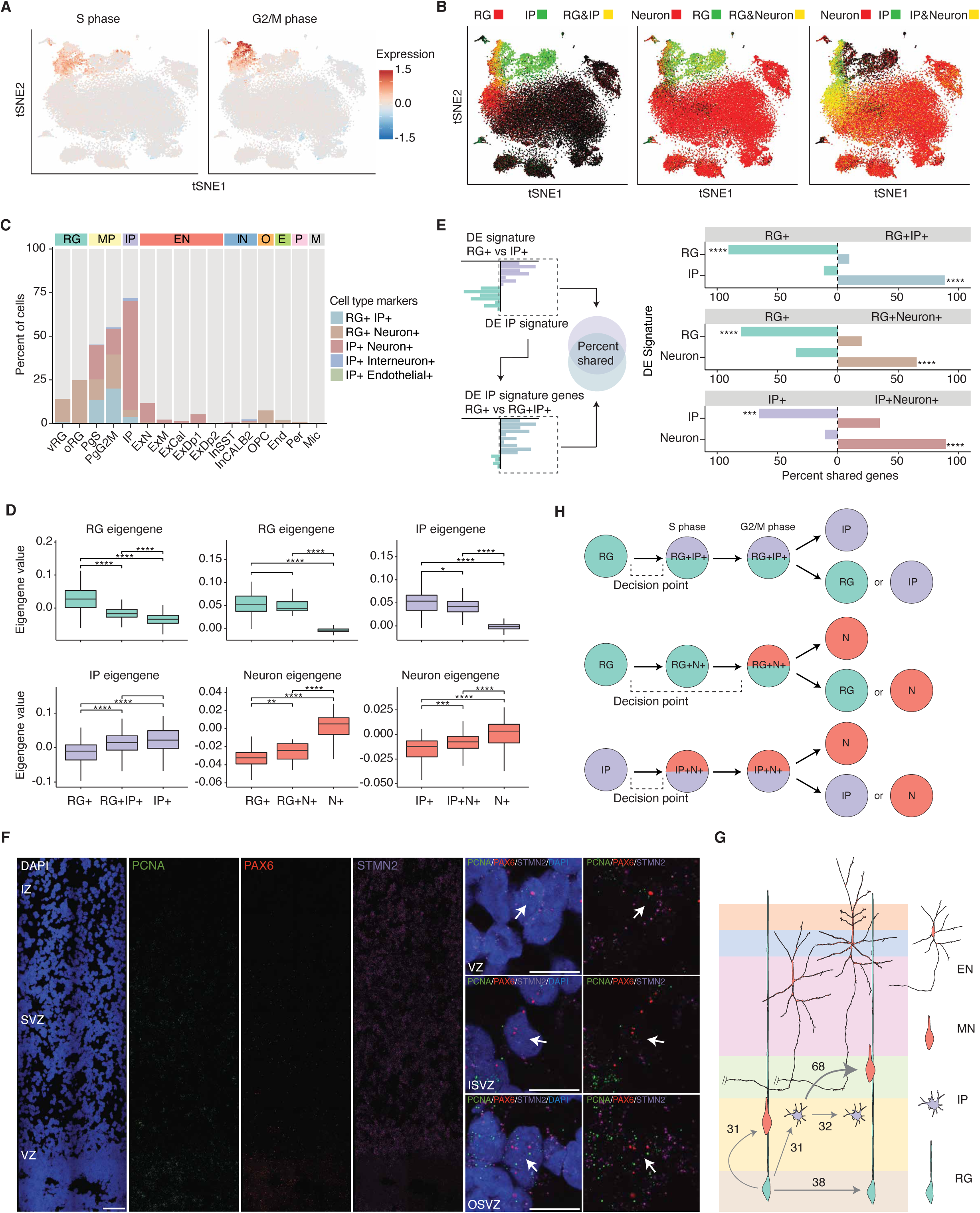
Dissecting the acquisition of a neuronal program. (A) tSNE colored by mean expression of cell cycle phase markers. (B) tSNE colored by co-expression of groups of canonical cell type markers. Yellow indicates co-expression. (C) Percent of cells in each Seurat cluster displaying co-expression of major cell type markers. (D) Mixed transcriptomic signatures of mixed marker cells in S-phase corresponding to expression of markers from multiple cell types. For the RG to IP comparison, RG and IP eigengenes were derived from differentially expressed genes between RG and IP cells, and similarly for the RG to Neuron comparison and IP to Neuron comparison. Boxplots: box indicates first and third quartiles; the whiskers extend from the box to the highest or lowest value that is within 1.5 * inter-quartile range of the box; and the line is the median. (E) Shared gene signatures between major cell types and mixed cell types. (F) RNA FISH of fetal cortex probed with the S-phase marker PCNA (green), the RG marker PAX6 (red), the neuron marker STMN2 (magenta), and stained with DAPI (blue). Panels on the right show high magnification single-plane confocal images of individual cells expressing all three markers. Scale bar = 100μm (left) or 10μm (right). (G) Quantification of relative amounts of mitotic progenitors undergoing different differentiation events. (H) Diagram of mixed cell type transcriptomic states that is characteristic of neurogenic differentiation trajectories in human neocortex. RG: radial glia; MP: mitotic progenitor; IP: intermediate progenitor; EN: excitatory neuron; IN: interneuron; O: oligodendrocyte precursor; E: endothelial cell; P: pericyte; M: microglia.

Therefore, as an alternative explanation, we hypothesized that we were identifying an intermediate or transition state: mitotically active cells in the early stages of neurogenesis, *i.e.* RG producing IP, RG producing neurons, and IP producing neurons. Consistent with this hypothesis, mixed marker cells progressing through different stages of the cell cycle consistently displayed transcriptomes comprised of multiple major cell types (**Fig. 3D** and fig. S11, A and B). By S-phase, RG+IP+ and IP+Neuron+ cells more closely resembled their presumed endpoint cell type, IP and neuron, respectively (**Fig. 3D** and fig S11B). The transcriptomic signature of RG+Neuron+ S-phase and G2/M phase cells was closer to RG, potentially reflecting the greater dissimilarity between RG and neurons (**Fig. 3D** and fig. S11, A and B). In addition, the mixed marker cells share a high percentage of the end point cell type signature, but the magnitude of expression of the cell type relevant signature genes is smaller than in cells in the fully differentiated cell clusters (**Fig. 3E** and fig. S11, C to F). Mixed marker cells not in S, G2, or M phase may represent cells starting to cycle and differentiate, consistent with findings in mice that some radial glial precursors also express neuronal marker genes of both deep and superficial layers, representing transcriptionally primed cells (*35*). Alternatively, these mixed marker cells may be newborn cells that still retain some transcripts of the mother cell type, as has been previously suggested in mice (*18, 35*).

To independently validate the existence of cells in these transition states, we performed RNA FISH, observing S-phase neural progenitors in the VZ expressing both PAX6 and STMN2, indicating an induction of a neuronal program in a single cell before its neurogenic division (**Fig. 3F**). Quantification of 872 cells across the germinative layers indicated that 6.1% (VZ), 5.8% (iSVZ) and 8.2% (oSVZ) of these cells co-express markers of RG and neurons, confirming our scRNA-seq data (see materials and methods). Given that mixed marker cells are indicative of a single cell transition state during early neurogenesis, we were able to quantify the relative proportions of progenitors undergoing distinct differentiation divisions (**Fig. 3G**), finding that RG produce roughly equal numbers of RG, IP, and neurons, but that IP produce approximately two times as many neuronal progeny as IP progeny. Taken together, these results indicate that during early neurogenesis: 1) Cell fate decisions occur prior to S-phase; 2) Differentiating “parent” cells not only express the few key transcription factors that drive cell fates, but express broad, mixed cell type transcriptomes; 3) Neural cell-type differentiation occurs on a continuum and involves transcriptomic transitions tied to cell cycle progression (**Fig. 3H**). An early cell fate decision point tied to cell cycle is consistent with previous work indicating cell fate decisions in neurogenesis are made in G1 (*36, 37*). However, previous models of asymmetric neurogenic divisions suggest that only a few key transcription factors of the “daughter” lineage are expressed in the asymmetrically dividing cell, whereas we observe early induction of more extensive cell type transcriptional programs (*36, 37*). Cells are producing transcriptomes representing two cell types from a single nucleus, prior to cytokinesis, demonstrating the usefulness of single-cell transcriptomics for furthering our understanding the transcriptional dynamics involved in early neural development (*38, 39*).

We next reasoned that we could begin to leverage these single cell data to uncover some of the cellular and molecular mechanisms driving human cortical evolution by determining whether specific cell types were enriched with genes showing human-specific expression trajectories (hSET) in bulk tissue (*40*) (see materials and methods). We observed the strongest enrichment of hSET genes in outer radial glia (oRG) and callosal neurons, an upper layer excitatory subtype (fig. S12A). This is notable, since both of these cell types represent core processes involved in both neocortical expansion (*16*) and the elaboration of extensive cortical connectivity in humans (*41*). Among the approximately 600 genes with oRG-enriched expression, we identified LYN, a Src tyrosine kinase previously implicated in neuronal polarization and AMPA signaling (*42, 43*), which had not been previously associated with this cell type. LYN also displayed VZ and oSVZ enrichment in a fetal LCM atlas (*30*) (fig. S12B). We used RNA FISH to further validate these observations in developing human brains, showing that LYN localized to the germinal zones and was specifically expressed in the VZ and oSVZ, as measured by RNA FISH (fig. S12C).

Using a similar logic, we reasoned that we could use this deep molecular atlas of developing human brain cell types to identify the developmental stages and cell types where mutations causing high risk for neuropsychiatric disease act, so as to provide a reference for understanding disease mechanisms and circuits. We first examined enrichment of high confidence autism spectrum disorder (ASD) risk genes, defined by harboring high risk likely protein-disrupting mutations (*44*) (**Fig. 4A** and fig. S13). The majority of ASD-risk genes were expressed in developing glutamatergic neurons, both deep and upper layer (**Fig. 4A**), consistent with previous studies (*45, 46*). However, at the individual gene level, there is substantial variability, and several genes are expressed in inhibitory neurons as well as excitatory neurons or progenitors (**Fig. 4A** and fig. S13). For example, MYT1L and AKAP9 display pan neuronal expression, whereas GRIN2B is glutamatergic subtype specific, and ILF2 is expressed in cycling progenitors (**Fig. 4A** and fig. S13). In adult, expression again concentrated in glutamatergic neurons, with some genes exhibiting more pan-neuronal expression patterns.

**Fig. 4.**
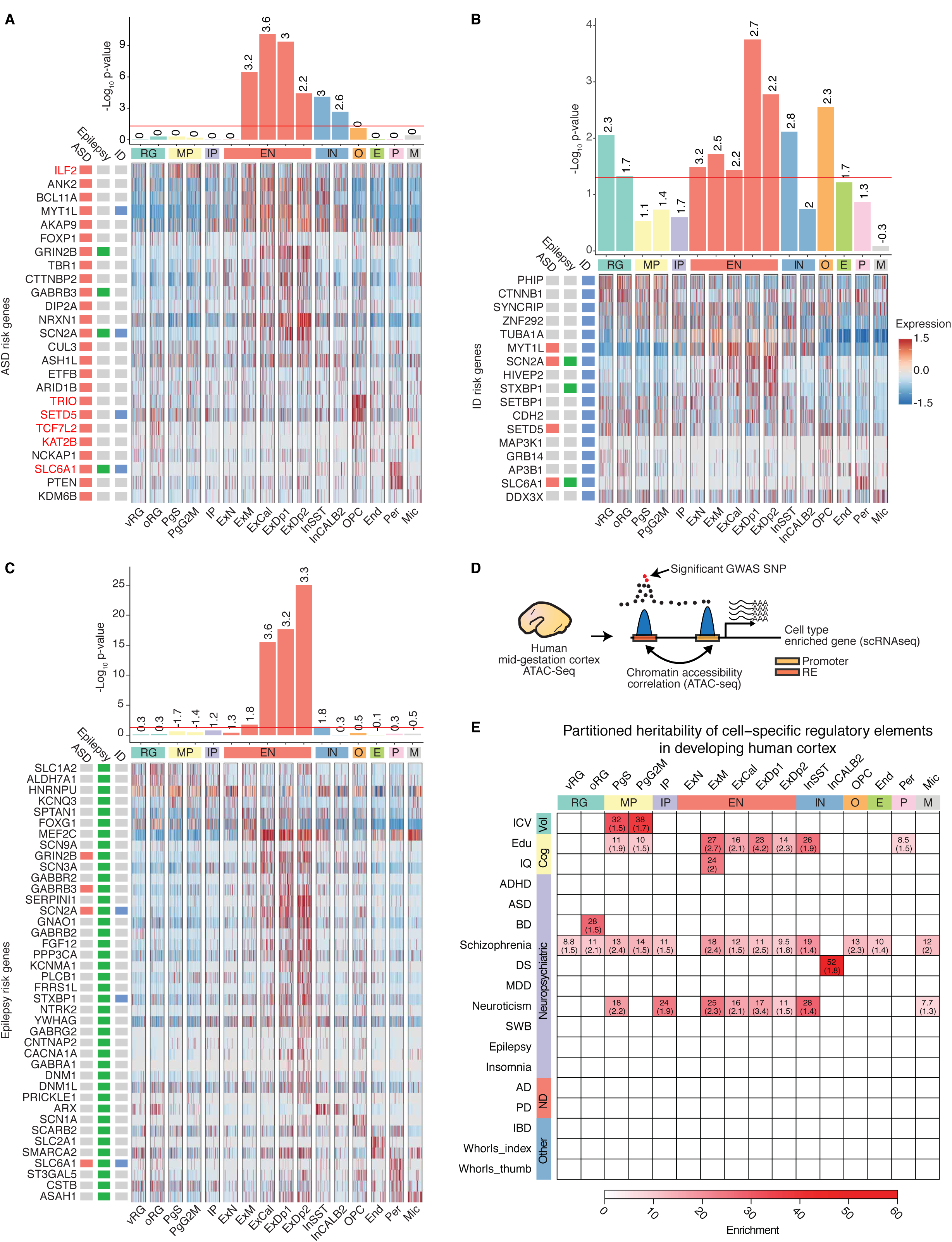
Cellular determinants of disease. (A to C) Cell type enrichment of ASD, ID, or epilepsy risk genes respectively. Heatmap: Expression of ASD risk genes is enriched in fetal glutamatergic neurons with some genes specifically expressed in other cell types. Red: gene is discussed in text. Bar graphs: numbers indicate log_2_ odds ratio, the red line indicates significance. Heatmaps: cells are ordered by cluster. RG: radial glia; MP: mitotic progenitor; IP: intermediate progenitor; EN: excitatory neuron; IN: interneuron; O: oligodendrocyte precursor; E: endothelial cell; P: pericyte; M: microglia. (D) Schematic showing the approach to identify regulatory elements (RE) for specific cell types and assess enrichment for specific brain traits. REs of genes enriched in specific cell types are identified by chromatin accessibility correlation between the promoter of the gene and other accessible peaks within 1Mb. The set of promoter and distal RE peaks are then tested for enrichment in SNPs associated with brain traits and neuropsychiatric disease using partitioned heritability by LD score regression. (E) Heatmap showing significant partitioned heritability enrichment for specific brain traits and neuropsychiatric disorders in different cell populations identified in developing human cortex. The color indicates the partitioned heritability enrichment and numbers are the FDR-corrected P-values. Only significant enrichments are shown in the heatmap, complete results are found in fig. S17. References for each GWAS are found in table S8.

In addition, our expanded atlas of cell types identified several genes that showed remarkably distinct patterns of extra-neuronal expression, including SLC6A1, which was enriched in pericytes, and TRIO, SETD5, TCF7L2, and KAT2B, which were enriched in oligodendrocyte precursors (**Fig. 4A** and fig. S13). These data implicate these cell types involved in maintenance of the blood brain barrier and the peri-neural environment for the first time in ASD risk. We then expanded this analysis to high confidence intellectual disability (ID) and epilepsy risk genes, (**Fig. 4, B and C**, and figs. S14 to S16). The majority of epilepsy-risk genes are also expressed in glutamatergic neurons in fetal and adult cortex (**Fig. 4C** and figs. S14 and S15). ID risk genes were also enriched in glutamatergic neurons, but interestingly also showed enrichment in RG, which was not observed with ASD or epilepsy. The impact on early progenitor types in ID relative to ASD and epilepsy is consistent with the more severe disease phenotype in ID (**Fig. 4B** and fig. S16). Taken together, these results demonstrate cell type specific expression of ASD, epilepsy, and ID risk genes by mid fetal development and provide a framework for the cellular and developmental context in which individual ASD, epilepsy, and ID genes should optimally be studied.

The majority of neuropsychiatric disease risk loci are found in the non-coding genome, where functional interpretation is hampered by limited knowledge of the genomic location and spatiotemporal activity of regulatory elements. Leveraging our cell type specific map of regulatory elements active in the human neocortex (**Fig. 4D** and table S7, see materials and methods), (*29*) we used a partitioned heritability approach based on LD score regression (*47*) to identify cell types enriched for variants influencing brain volume, cognition, or causing risk for neuropsychiatric disease (*48-63*). We found that variants influencing adult intracranial volume were specifically enriched in the regulatory elements of cycling progenitors (PgS, PgG2M), pinpointing a specific cell type and state likely associated with neural progenitor expansion (**Fig. 4E** and fig. S17). Importantly, by connecting causal genetic drivers to specific genes within a specific cell type, this not only identifies putative cell-type specific mechanisms involved in cortical expansion, but provides further support for the radial unit hypothesis of cortical expansion on the human lineage (*16, 64*).

In contrast, common genetic variants influencing general cognition (IQ) were enriched in cycling neural progenitors, cortical plate glutamatergic neurons, SST inhibitory neurons, and intriguingly in pericytes (**Fig. 4E** and fig. S17). A less powered IQ genome-wide association study (GWAS) also found enrichment in maturing cortical plate glutamatergic neurons, but not in other cell types (**Fig. 4E** and fig. S17). Variants causing risk for schizophrenia were enriched in multiple cell types, including neural progenitors, glutamatergic neurons, interneurons, oligodendrocyte precursors and microglia **(Fig. 4E** and fig. S17). A recent study, using a partitioned heritability approach, found enrichment for schizophrenia variants in adult cortical glutamatergic neurons and cortical interneurons, consistent with bulk tissue analysis (*65*), but was unable to assess enrichment in human fetal cortical cell types given a lack of available data (*66*). Thus, our results now also implicate neural progenitors, oligodendrocyte precursors and fetal microglia in schizophrenia, highlighting the importance of generating single-cell resources from multiple time periods and brain regions. We did not observe enrichment of IBD or finger whorl variants (**Fig. 4E** and fig. S17) in the regulatory elements of any of the cortical derived cell types, supporting the cell type specificity of gene regulation. These results highlight how combining DNA accessibility profiling and single-cell sequencing can facilitate interpretation of the function of variants influencing brain structure and function.

This resource of transcriptomic profiles of 40,000 single cells in human fetal cortex demonstrates the utility of single-cell analysis for characterizing human neurogenesis, identifying novel cell type regulatory mechanisms, and for understanding the cellular basis of brain phenotypes with neurodevelopmental origins. By expanding the publicly available number of human fetal brain single-cell transcriptomes by an order of magnitude, these data provide more complete cell type mRNA transcript profiles, and permit discovery and characterization of rare cell types and states (*18-22*). These data implicate early decision points in cell fate trajectories that are pre S-phase, leading to transcriptomically mixed cell states, and highlighting the importance of sampling single cells at a massive scale to capture rare types and states. The transition state dynamics during early neurogenesis show that cell type differentiation is on a gradual continuum and involves transcriptomic transitions tied to cell cycle progression.

We also identify novel cell type enrichment of regulatory genes and elements and highlight how this can be used to identify critical cell types in monogenic disorders (e.g. ZFHX4 and 8q21.11 deletion), as well as in ASD, expanding the implicated cell landscape in this disorder to include inhibitory neurons and non-neural cells in addition to glutamatergic neurons. The importance of cell type specific gene regulation underscores the importance of expanding single-cell taxonomies in the future to include single-cell epigenetic analysis to supplement the rich information contained in scRNA-seq catalogs as recently demonstrated (*67*). Lastly, we show that genes with human specific expression patterns act preferentially in oRG and upper cortical layer neurons, which is consistent with the expansion of these zones during brain evolution. These data provide a molecular context for cortical expansion and increased cortical-cortical connectivity in humans, and extend our understanding of developmental dynamics and the origin of neuropsychiatric disease risk in human neocortex.

## Acknowledgements

Drop-seq libraries were sequenced by the UCLA BSCRC, RNA-seq libraries were prepared and sequenced by the UCLA Neuroscience Genomics Core, and fetal tissue was collected from the UCLA CFAR (5P30 AI028697). Finger whorl GWAS data were kindly provided by Sarah Medland. We thank members of the Geschwind lab for helpful discussions and critical reading of the manuscript. **Funding:** This work was supported by NIH grants to D.H.G. (1U01 MH105991, 5R01 MH081754, 5R01 MH100027, 1R01MH110927 and 1U01MH116489) and the Allen Distinguished Investigator Program to DHG, the California Institute for Regenerative Medicine (CIRM)-BSCRC Training Grant (TG2-01169) to L.T.U., K.P. was supported by NIH (P01GM099134) and the Allen Distinguished Investigator Program and the Paul G. Allen Frontiers Group, J. L. by the UCLA Tumor Cell Biology Training Program (USHHS Ruth L. Kirschstein Institutional National Research Service Award # T32 CA009056), and S. S. by the UCLA Broad Stem Cell Research Center – Rose Hills Foundation Training Award. **Author contributions:** L.T.U, J.S., and D.H.G. designed the study. DHG oversaw all of the analyses and supervised the study, with help from K.P., and B.L. L.T.U, J.L., C.K.O., and D.L., performed the single-cell experiments. D.P. and J.L.S. processed and managed the data. D.P., L.T.U., A.G.E., and W.C., analyzed the data, with input from S.S. L.T.U., D.L, and C.K.V. performed the validation experiments. T.H. and J.K.L performed CNV analyses.. D.P., L.T.U., and D.H.G. wrote and prepared the manuscript, with feedback from all authors. **Competing interests:** The authors declare no competing interests. **Data and materials availability:** The transcriptomic dataset is in the process of being deposited at dbGaP.

## Supplementary Materials

### Supplementary Materials

Materials and Methods

Figures S1-S17

Tables S1-S8

References (68-80)

## References and Notes

1. J. C. Silbereis, S. Pochareddy, Y. Zhu, M. Li, N. Sestan, The Cellular and Molecular Landscapes of the Developing Human Central Nervous System. Neuron 89, 248–268 (2016).

2. E. Lein, L. E. Borm, S. Linnarsson, The promise of spatial transcriptomics for neuroscience in the era of molecular cell typing. Science 358, 64–69 (2017).

3. C. F. Stevens, Neuronal diversity: too many cell types for comfort? Curr Biol 8, R708–710 (1998).

4. C. Dehay, H. Kennedy, K. S. Kosik, The outer subventricular zone and primate-specific cortical complexification. Neuron 85, 683–694 (2015).

5. E. Z. Macosko et al., Highly Parallel Genome-wide Expression Profiling of Individual Cells Using Nanoliter Droplets. Cell 161, 1202–1214 (2015).

6. A. Saunders et al., Molecular Diversity and Specializations among the Cells of the Adult Mouse Brain. Cell 174, 1015–1030 e1016 (2018).

7. K. Shekhar et al., Comprehensive Classification of Retinal Bipolar Neurons by Single-Cell Transcriptomics. Cell 166, 1308–1323 e1330 (2016).

8. B. Tasic et al., Adult mouse cortical cell taxonomy revealed by single cell transcriptomics. Nat Neurosci 19, 335–346 (2016).

9. A. Zeisel et al., Molecular Architecture of the Mouse Nervous System. Cell 174, 999–1014 e1022 (2018).

10. S. Hrvatin et al., Single-cell analysis of experience-dependent transcriptomic states in the mouse visual cortex. Nat Neurosci 21, 120–129 (2018).

11. J. R. Ecker et al., The BRAIN Initiative Cell Census Consortium: Lessons Learned toward Generating a Comprehensive Brain Cell Atlas. Neuron 96, 542– 557 (2017).

12. B. I. Bae, D. Jayaraman, C. A. Walsh, Genetic Changes Shaping the Human Brain. Developmental cell 32, 423–434 (2015).

13. D. H. Geschwind, P. Rakic, Cortical evolution: judge the brain by its cover. Neuron 80, 633–647 (2013).

14. T. Sun, R. F. Hevner, Growth and folding of the mammalian cerebral cortex: from molecules to malformations. Nat Rev Neurosci 15, 217–232 (2014).

15. E. Taverna, M. Gotz, W. B. Huttner, The cell biology of neurogenesis: toward an understanding of the development and evolution of the neocortex. Annual review of cell and developmental biology 30, 465–502 (2014).

16. J. H. Lui, D. V. Hansen, A. R. Kriegstein, Development and evolution of the human neocortex. Cell 146, 18–36 (2011).

17. Z. Molnar, Evolution of cerebral cortical development. Brain Behav Evol 78, 94– 107 (2011).

18. S. Zhong et al., A single-cell RNA-seq survey of the developmental landscape of the human prefrontal cortex. Nature 555, 524–528 (2018).

19. T. J. Nowakowski et al., Spatiotemporal gene expression trajectories reveal developmental hierarchies of the human cortex. Science 358, 1318–1323 (2017).

20. A. A. Pollen et al., Molecular identity of human outer radial glia during cortical development. Cell 163, 55–67 (2015).

21. S. J. Liu et al., Single-cell analysis of long non-coding RNAs in the developing human neocortex. Genome Biol 17, 67 (2016).

22. X. Fan et al., Spatial transcriptomic survey of human embryonic cerebral cortex by single-cell RNA-seq analysis. Cell Res 28, 730–745 (2018).

23. M. J. Gandal, V. Leppa, H. Won, N. N. Parikshak, D. H. Geschwind, The road to precision psychiatry: translating genetics into disease mechanisms. Nat Neurosci 19, 1397–1407 (2016).

24. L. de la Torre-Ubieta, H. Won, J. L. Stein, D. H. Geschwind, Advancing the understanding of autism disease mechanisms through genetics. Nat Med 22, 345–361 (2016).

25. A. Butler, P. Hoffman, P. Smibert, E. Papalexi, R. Satija, Integrating single-cell transcriptomic data across different conditions, technologies, and species. Nat Biotechnol 36, 411–420 (2018).

26. X. Qiu et al., Single-cell mRNA quantification and differential analysis with Census. Nat Methods 14, 309–315 (2017).

27. C. Trapnell et al., The dynamics and regulators of cell fate decisions are revealed by pseudotemporal ordering of single cells. Nat Biotechnol 32, 381–386 (2014).

28. A. T. Lun, K. Bach, J. C. Marioni, Pooling across cells to normalize single-cell RNA sequencing data with many zero counts. Genome Biol 17, 75 (2016).

29. L. de la Torre-Ubieta et al., The Dynamic Landscape of Open Chromatin during Human Cortical Neurogenesis. Cell 172, 289–304 e218 (2018).

30. J. A. Miller et al., Transcriptional landscape of the prenatal human brain. Nature 508, 199–206 (2014).

31. A. Hoerder-Suabedissen, Z. Molnar, Development, evolution and pathology of neocortical subplate neurons. Nat Rev Neurosci 16, 133–146 (2015).

32. F. M. Oeschger et al., Gene expression analysis of the embryonic subplate. Cereb Cortex 22, 1343–1359 (2012).

33. M. Palomares et al., Characterization of a 8q21.11 microdeletion syndrome associated with intellectual disability and a recognizable phenotype. Am J Hum Genet 89, 295–301 (2011).

34. D. V. Hansen, J. H. Lui, P. R. Parker, A. R. Kriegstein, Neurogenic radial glia in the outer subventricular zone of human neocortex. Nature 464, 554–561 (2010).

35. S. K. Zahr et al., A Translational Repression Complex in Developing Mammalian Neural Stem Cells that Regulates Neuronal Specification. Neuron 97, 520–537 e526 (2018).

36. S. Dalton, Linking the Cell Cycle to Cell Fate Decisions. Trends Cell Biol 25, 592–600 (2015).

37. B. Pfeuty, A computational model for the coordination of neural progenitor selfrenewal and differentiation through Hes1 dynamics. Development 142, 477–485 (2015).

38. E. B. Dewey, D. T. Taylor, C. A. Johnston, Cell Fate Decision Making through Oriented Cell Division. J Dev Biol 3, 129–157 (2015).

39. V. Bertrand, O. Hobert, Lineage programming: navigating through transient regulatory states via binary decisions. Curr Opin Genet Dev 20, 362–368 (2010).

40. T. E. Bakken et al., A comprehensive transcriptional map of primate brain development. Nature 535, 367–375 (2016).

41. R. M. Fame, J. L. MacDonald, J. D. Macklis, Development, specification, and diversity of callosal projection neurons. Trends Neurosci 34, 41–50 (2011).

42. T. Namba et al., Pioneering axons regulate neuronal polarization in the developing cerebral cortex. Neuron 81, 814–829 (2014).

43. T. Hayashi, H. Umemori, M. Mishina, T. Yamamoto, The AMPA receptor interacts with and signals through the protein tyrosine kinase Lyn. Nature 397, 72–76 (1999).

44. S. J. Sanders et al., Insights into Autism Spectrum Disorder Genomic Architecture and Biology from 71 Risk Loci. Neuron 87, 1215–1233 (2015).

45. N. N. Parikshak et al., Integrative functional genomic analyses implicate specific molecular pathways and circuits in autism. Cell 155, 1008–1021 (2013).

46. T. Bourgeron, From the genetic architecture to synaptic plasticity in autism spectrum disorder. Nat Rev Neurosci 16, 551–563 (2015).

47. H. K. Finucane et al., Partitioning heritability by functional annotation using genome-wide association summary statistics. Nat Genet 47, 1228–1235 (2015).

48. N. R. Wray et al., Genome-wide association analyses identify 44 risk variants and refine the genetic architecture of major depression. Nat Genet 50, 668–681 (2018).

49. A. F. Pardinas et al., Common schizophrenia alleles are enriched in mutation-intolerant genes and in regions under strong background selection. Nat Genet 50, 381–389 (2018).

50. H. H. Adams et al., Novel genetic loci underlying human intracranial volume identified through genome-wide association. Nat Neurosci 19, 1569–1582 (2016).

51. International League Against Epilepsy Consortium on Complex Epilepsies, Genetic determinants of common epilepsies: a meta-analysis of genome-wide association studies. Lancet Neurol 13, 893–903 (2014).

52. J. Grove et al., Common risk variants identified in autism spectrum disorder. bioRxiv, (2017).

53. S. Sniekers et al., Genome-wide association meta-analysis of 78,308 individuals identifies new loci and genes influencing human intelligence. Nat Genet 49, 1107–1112 (2017).

54. D. Demontis et al., Discovery Of The First Genome-Wide Significant Risk Loci For ADHD. bioRxiv, (2017).

55. Psychiatric Gwas Consortium Bipolar Disorder Working Group, Large-scale genome-wide association analysis of bipolar disorder identifies a new susceptibility locus near ODZ4. Nat Genet 43, 977–983 (2011).

56. M. Luciano et al., Association analysis in over 329,000 individuals identifies 116 independent variants influencing neuroticism. Nat Genet 50, 6–11 (2018).

57. A. R. Hammerschlag et al., Genome-wide association analysis of insomnia complaints identifies risk genes and genetic overlap with psychiatric and metabolic traits. Nat Genet 49, 1584–1592 (2017).

58. A. Okbay et al., Genetic variants associated with subjective well-being, depressive symptoms, and neuroticism identified through genome-wide analyses. Nat Genet 48, 624–633 (2016).

59. A. Okbay et al., Genome-wide association study identifies 74 loci associated with educational attainment. Nature 533, 539–542 (2016).

60. J. C. Lambert et al., Meta-analysis of 74,046 individuals identifies 11 new susceptibility loci for Alzheimer’s disease. Nat Genet 45, 1452–1458 (2013).

61. Y. Y. Ho et al., Common Genetic Variants Influence Whorls in Fingerprint Patterns. J Invest Dermatol 136, 859–862 (2016).

62. L. Jostins et al., Host-microbe interactions have shaped the genetic architecture of inflammatory bowel disease. Nature 491, 119–124 (2012).

63. M. A. Nalls et al., Large-scale meta-analysis of genome-wide association data identifies six new risk loci for Parkinson’s disease. Nat Genet 46, 989–993 (2014).

64. P. Rakic, A small step for the cell, a giant leap for mankind: a hypothesis of neocortical expansion during evolution. Trends Neurosci 18, 383–388 (1995).

65. S. Horvath, K. Mirnics, Schizophrenia as a disorder of molecular pathways. Biol Psychiatry 77, 22–28 (2015).

66. N. G. Skene et al., Genetic identification of brain cell types underlying schizophrenia. Nat Genet 50, 825–833 (2018).

67. C. Luo et al., Single-cell methylomes identify neuronal subtypes and regulatory elements in mammalian cortex. Science 357, 600–604 (2017).

68. N. J. Cooper et al., Detection and correction of artefacts in estimation of rare copy number variants and analysis of rare deletions in type 1 diabetes. Hum Mol Genet 24, 1774–1790 (2015).

69. A. Dobin et al., STAR: ultrafast universal RNA-seq aligner. Bioinformatics 29, 15– 21 (2013).

70. H. Li et al., The Sequence Alignment/Map format and SAMtools. Bioinformatics 25, 2078–2079 (2009).

71. S. Anders, P. T. Pyl, W. Huber, HTSeq--a Python framework to work with high-throughput sequencing data. Bioinformatics 31, 166–169 (2015).

72. C. Hennig, Cluster-wise assessment of cluster stability. Comput Stat Data An 52, 258–271 (2007).

73. J. Reimand et al., g:Profiler-a web server for functional interpretation of gene lists (2016 update). Nucleic Acids Res 44, W83–89 (2016).

74. H. M. Zhang et al., AnimalTFDB 2.0: a resource for expression, prediction and functional study of animal transcription factors. Nucleic Acids Res 43, D76–81 (2015).

75. J. D. Buenrostro, P. G. Giresi, L. C. Zaba, H. Y. Chang, W. J. Greenleaf, Transposition of native chromatin for fast and sensitive epigenomic profiling of open chromatin, DNA-binding proteins and nucleosome position. Nat Methods 10, 1213–1218 (2013).

76. B. B. Lake et al., Integrative single-cell analysis of transcriptional and epigenetic states in the human adult brain. Nat Biotechnol 36, 70–80 (2018).

77. A. Rauch et al., Range of genetic mutations associated with severe non-syndromic sporadic intellectual disability: an exome sequencing study. Lancet 380, 1674–1682 (2012).

78. J. de Ligt et al., Diagnostic exome sequencing in persons with severe intellectual disability. N Engl J Med 367, 1921–1929 (2012).

79. F. Wang et al., RNAscope: a novel in situ RNA analysis platform for formalin-fixed, paraffin-embedded tissues. J Mol Diagn 14, 22–29 (2012).

80. C. Gawad, W. Koh, S. R. Quake, Single-cell genome sequencing: current state of the science. Nat Rev Genet 17, 175–188 (2016).

